# Accelerated Measurement of Chemical Exchange Saturation Transfer by Accordion NMR Spectroscopy

**DOI:** 10.64898/2026.07.01.735851

**Authors:** Göran Carlström, Theresa Höfurthner, Mikael Akke

## Abstract

Chemical exchange saturation transfer (CEST) has become an indispensable NMR method to characterize slow exchange affecting biomacromolecules, especially for cases involving exchange between a major state and a minor state, the latter of which is often invisible in the spectrum. The CEST method is based on successive irradiation of selective regions of the NMR spectrum using a weak radiofrequency field, *B*_1_, while observing the effect on the visible major state when the *B*_1_ field saturates the invisible minor state. The need for selective saturation of narrow spectral regions has to date required acquisition of many tens of two-dimensional CEST spectra to sample the entire spectrum with sufficient resolution. Here we present the ACCEST method which measures an entire CEST profile from a single two-dimensional accordion-CEST spectrum plus a reference spectrum. ACCEST is based on the concept of accordion spectroscopy, where in the present implementation the carrier frequency of the weak saturating *B*_1_ field is stepped in synchrony with the dwell-time incrementation in the indirect dimension of the two-dimensional spectrum. We benchmarked ACCEST against conventional CEST, resulting in excellent agreement for both backbone ^15^N and methyl ^13^C CEST profiles. ACCEST offers substantial time savings that scale linearly with the number of spectra required in the corresponding conventional CEST experiment. Thus, ACCEST can dramatically speed up lengthy serial experiments, such as ligand titrations or temperature-dependent studies, and enable studies of non-equilibrium systems or samples with limited lifetimes.

---

NMR spectroscopy is a powerful tool for studying conformational dynamics in biomolecules, providing insights into transient states that are crucial for biological function. By capturing motions on timescales ranging from microseconds to milliseconds, NMR enables the characterization of conformational transitions that underlie key processes such as enzyme catalysis, protein folding, molecular recognition, and allostery.^1-5^ One particularly effective NMR method for probing slow conformational exchange (milliseconds to seconds) is chemical exchange saturation transfer (CEST), which exploits the transfer of magnetization between exchanging states to detect low-populated conformers with high sensitivity.^6, 7^ CEST has been instrumental in uncovering such ‘invisible’ high-energy states of proteins and nucleic acids, shedding light on mechanisms involved in allosteric regulation, ligand binding, and disease-related protein misfolding.^8-12^

In a typical CEST experiment, a weak radiofrequency (RF) field (*B*_1_) is applied at the resonance frequency of the invisible minor state, causing continuous saturation. Due to exchange, saturation transfer to the observable major state results in a measurable reduction in its signal intensity. By systematically varying the saturation frequency and measuring the resulting signal changes, a CEST profile is generated, revealing information about exchange rates, populations, and chemical shifts of the minor states. The need to acquire one two-dimensional (2D) spectrum for each saturation frequency of the CEST profile makes the entire experiment quite time consuming. Depending on the static magnetic field strength and the frequency difference between the major and minor states, the number of spectra often ranges from many tens up to several hundreds.

Recent developments of CEST have focused on accelerating data acquisition while maintaining sensitivity to low-populated conformational states. One key advancement is the implementation of multi-frequency excitation methods^13, 14^ that simultaneously probe multiple frequencies in a single spectrum. However, this approach typically still requires acquisition of tens of spectra to probe the entire frequency range. Other developments serve to acquire several CEST experiments on different nuclei in rapid succession,^15^ which expedites data acquisition in those cases where it is desired to probe the exchange process of several types of atoms. These innovations aim to shorten the overall experiment time, which is of prime interest for the purpose of optimizing usage of expensive instrumentation and enabling studies of exchange processes under nonequilibrium conditions, such as amyloid-forming or other aggregating systems, or in samples of limited stability, such as living cells.

Here we present the ACCEST experiment, which produces an entire CEST profile from only two 2D spectra: one accordion-CEST spectrum and one reference spectrum, thereby delivering dramatic time savings compared to previous approaches. The ACCEST method is a novel variant of accordion spectroscopy,^16-19^ where the carrier frequency of the *B*_1_ field is changed synchronously with the incrementation of the indirect evolution period (*t*_1_) of the two-dimensional spectrum. ACCEST is quite general and can be applied to a wide range of nuclei, coherence selections, and isotope labeling schemes.

## RESULTS

Below, we first describe the underlying theory of ACCEST and then benchmark it against conventional CEST using the G48A variant of the Fyn SH3 domain, a protein frequently used to validate new NMR relaxation methods.^20^ We recorded ACCEST experiments at 10° C on a 1 mM sample of the G48A Fyn SH3 domain, which undergoes folding-unfolding exchange dynamics under these conditions.^20^ This sample was uniformly isotope-enriched with ^15^N and selectively labeled with ^13^CH_3_ methyl groups in the δ positions of Ile and Leu and the γ positions of Val against an otherwise uniformly ^12^C/^2^H-labeled background.

### Measuring CEST profiles from only two spectra

The ACCEST pulse sequences are essentially identical to the original CEST versions,^7, 20^ except that the variable saturation frequency is accordion-encoded into the indirect dimension (*t*_1_) of the two-dimensional spectrum. This modification can easily be implemented in any existing CEST pulse sequence. In the resulting ACCEST spectrum, the *t*_1_ data column (interferogram) exhibits a decrease in intensity for those *t*_1_ points that correspond to the *B*_1_ carrier frequencies (*ω*_1_) matching the resonance frequencies of the minor or major states (Fig. 1).

**Figure 1.**
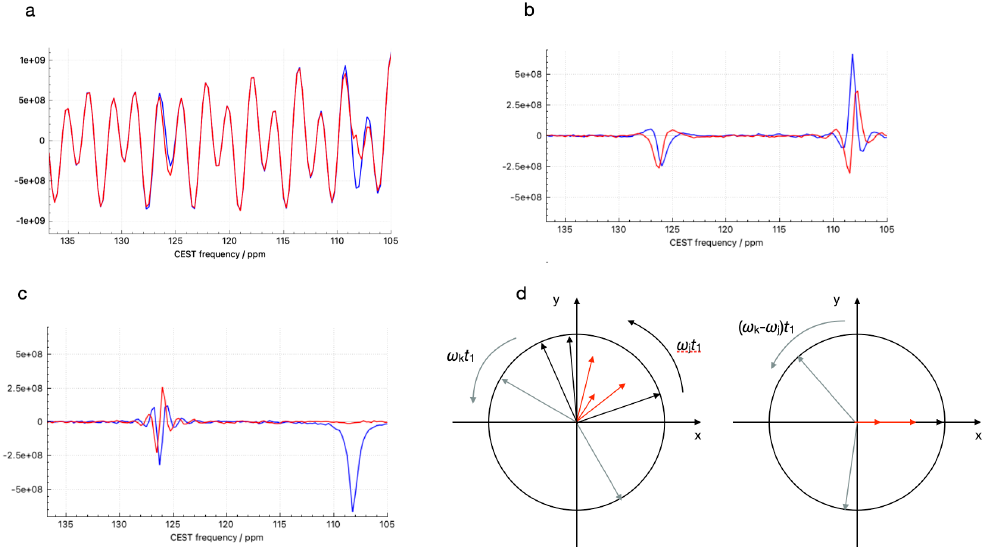
Schematic representation of the principles behind ACCEST. (a) Example *t*_1_ data (real part of the interferograms) containing two resonances. The reference data are shown in blue and the ACCEST data in red. Note that the time co-ordinate of the x-axis has been reversed to have the CEST frequency to increase towards the left, as is usual in NMR spectra. The two data sets perfectly superimpose, with the red covering the blue, except for those *t*_1_ data points where the carrier frequency (*ω*_1_) of the *B*_1_ field matches the resonance frequency of either a major or minor state. For these cases, the resonance of the major state will experience a reduction in signal intensity. (b) The difference data, Δ*S*, obtained by subtracting the reference data, *S*_ref_, from the raw ACCEST data, *S*_CEST_. (c) The frequency-shifted difference data, 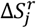. Note that this data representation is directly related to the CEST profile recorded by conventional means. In panels b and c, the real part of the signal is shown in blue, and the imaginary part in red. (d) Vector diagram illustrating the data shown in panels a–c. The phase of the evolving magnetization identifies which of the resonances contributing to a given interferogram is affected by the *B*_1_ field. In the diagram to the left, corresponding to panel b, the two resonances are indicated by black/red (resonance 1) and gray (resonance 2) vectors, where red indicates the time points where the major state of resonance 1 is affected by saturation. For clarity, only the first two *t*_1_ data points are displayed for resonance 2 (grey). Shifting the rotating frame frequency to the frequency of resonance 1, gives the vector diagram to the right, which corresponds to panel c. Here the magnetization from resonance 1 stays along the real axis, whereas the other resonance now rotates with a frequency *Ω*_k_-*Ω* _j_ and will also involve an imaginary signal. Relaxation effects and the effect of the irradiation on resonance 2 have been neglected for clarity.

The ACCEST data set can be described as follows

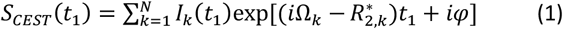

where *k* denotes the *N* signals present in a given *t*_1_ column, *I*_k_(*t*_1_), *Ω*_k_, and 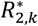 denote the intensity, the resonance offset and the effective transverse relaxation rate constant of signal *k*, respectively, and *φ* indicates a phase offset, which from here on is assumed to be zero (without loss of generality). The variation of the peak intensity with *ω*_1_ is indicated by the explicit time dependence, *I*_k_(*t*_1_). A reference experiment is performed with the *B*_1_ field applied far off-resonance, yielding the signal

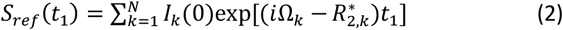

where the intensity *I*_k_(0) is constant. Figure 1a provides an example representation of *S*_CEST_ and *S*_ref_, showing the effect on *I*_k_(*t*_1_) when *ω* _1_ matches the resonance frequencies of the signals contributing to the interferogram.

Following subtraction of *S*_ref_ from *S*_CEST_, we obtain

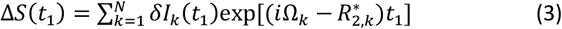

The intensities *δI*_k_(*t*_1_) = *I*_k_(*t*_1_) – *I*_k_(0) of Δ*S*(*t*) are zero at all *t*_1_ points except those where *ω*_1_ matches the resonance frequency of either a major (ground) state or a minor (high-energy) state (Fig. 1b). *δI*_k_(*t*_1_) will tend towards –*I*_k_(0) for the *t*_1_ points where *ω*_1_ matches the resonance frequency of either a major or minor state. Thus, *δI*_k_(*t*_1_) is directly related to the CEST profile, *C*_k_(*ω*_1_), from a conventional CEST experiment. *C*_k_(*ω*_1_) can be obtained as *C*_k_(*ω*_1_) = *δI*_k_(*ω*_1_(*t*_1_))/*I*_k_(0) + 1, where the linear mapping from *t*_1_ to *ω*_1_ is explicitly indicated. In the case where there is more than one signal in the *t*_1_ column, the resonance frequency of a given signal, denoted the signal of interest, is identified by comparing the phase evolution of the non-zero data points of Δ*S*(*t*_1_) with the phase evolution of each signal in the reference *t*_1_ column and finding the matching one. In practice, this identification is conveniently done by shifting the rotating-frame frequency to match the offset *Ω*_*j*_ of the signal of interest (Fig. 1c, d). Thus, we generate a frequency-shifted difference signal, Δ*S*^r^_*j*_(*t*_1_), in which both ‘auto’ and ‘CEST’ peaks of the signal of interest consist of a purely real component (while its imaginary component is zero):

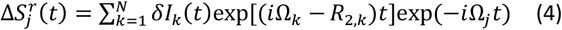

Signals in Δ*S*^r^_*j*_(*t*_1_) from other resonances will now evolve with a frequency *Ω*_k_–*Ω*_j_. If there is only a single exchanging signal in the *t*_1_ column, then the rotating frame transformation recovers the CEST profile obtained from a conventional experiment, showing both the ‘auto’ and ‘CEST’ dips as peaks in the real part of the complex signal, whereas the imaginary part is zero; see Fig. 1c, d. (If there is only a single non-exchanging signal, then naturally only the auto dip appears.) If there are multiple signals in the *t*_1_ column, then the imaginary channel of 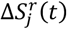 will include non-zero signals from spins *k* ≠ *j* (Fig. 1c, d). This feature resolves the issue when a CEST peak from a minor state of one residue overlaps the *t*_1_ points of a peak (CEST or auto) of another residue, so long as the major forms of the two residues have different resonance frequencies. Just as in a conventional CEST experiment, overlap between resonances of two or more major forms poses a problem. Note that the reference data set provides information on *I*_k_(0), *Ω*_k_, and 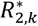 of all resonances in the *t*_1_ column and this prior knowledge thus makes it possible to subtract the overlapping signal component to reveal the CEST peak of interest. In practice, we perform these operations in an iterative procedure, see flowchart in SI Fig. 1, where the CEST profiles of all spins *j* and *k* are analyzed in series. The procedure converges rather fast, and normally only 2–3 iterations are required. An example of the procedure is shown in Fig. 2 for three signals, partly overlapping in the ^1^H dimension, in the ^15^N ACCEST 2D spectrum of Fyn SH3. To analyze the ACCEST data and reconstruct the CEST profile, we used DSURE,^21^ a fast and efficient algorithm for reconstructing the signals, but any method providing the parameters *I*_k_(0), *Ω*_k_, and 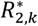 can be used.

**Figure 2.**
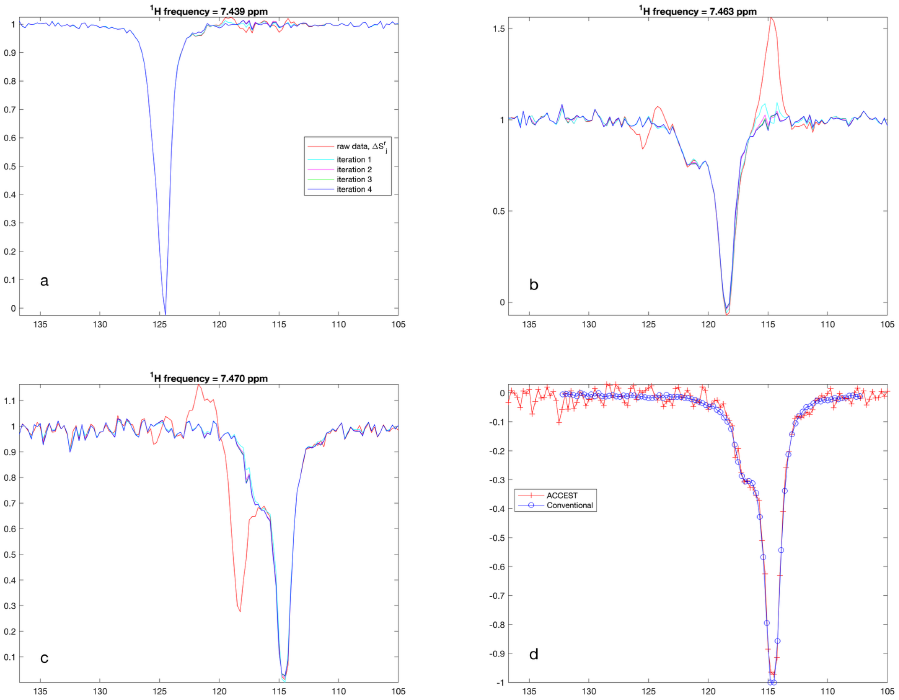
Results from a simultaneous and iterative analysis of three resonances appearing in the same *t*_1_ interferogram of an ^15^N ACCEST dataset. Four iterations were performed, with the CEST profile resulting after each iteration shown: red, raw data; cyan, magenta, green, and blue correspond to iterations 1–4, respectively. (a) Residue Arg60, δ(^1^H) = 7.439 ppm. (b) Residue Leu55, δ(^1^H) = 7.463 ppm. (c) Residue Tyr54, δ(^1^H) = 7.470 ppm. (d) Comparison of the CEST profiles obtained for residue Tyr54 with ^15^N chemical shift of 114.6 ppm using ACCEST (red) or conventional CEST (blue). The number of scans NS = 8 for both experiments.

The decay caused by transverse relaxation and magnetic inhomogeneity during the evolution period affects the sensitivity of the ACCEST experiment to varying degrees, depending on which *t*_1_ points correspond to the *B*_1_ frequencies matching the minor-state resonance. To minimize the longest sampled *t*_1_-value, we oversampled the data in the indirect dimension by using a spectral width two or three times wider than what is needed to cover region of interest. To weight each *t*_1_ point equally, we multiplied the obtained CEST profile by a factor 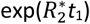. Another option is to implement a constant-time (CT) evolution period, which yields an essentially non-decaying interferogram with intensities reduced by a factor 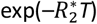, where *T* is the total length of the evolution period. Thus, the intensity in the ACCEST experiment is reduced by 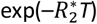 compared to that of a conventional CEST experiment, whereas the non-CT version of ACCEST is scaled by approximately exp(– 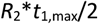 on average. The estimated error in the CEST peak intensity is obtained directly from the root-mean-square intensity calculated over empty regions of the imaginary component of Δ*S*^r^.

### ACCEST accurately reproduces results from conventional CEST while substantially shortening experiment time

Figure 3 shows comparisons of ^15^N and ^13^C methyl CEST profiles obtained from a conventional experiment and an ACCEST experiment, demonstrating excellent agreement. The duration of the ACCEST experiment is considerably shorter than alternative CEST approaches. In the present example (Fig. 3), the ^15^N and ^13^C ACCEST experiments each took 3 hours in total, including the reference experiment, whereas the conventional ^15^N and ^13^C CEST experiments were acquired in 60 and 30 hours, respectively. Thus, the ACCEST approach shortened the experiment time by a factor of 10 to 20 in this case.

**Figure 3.**
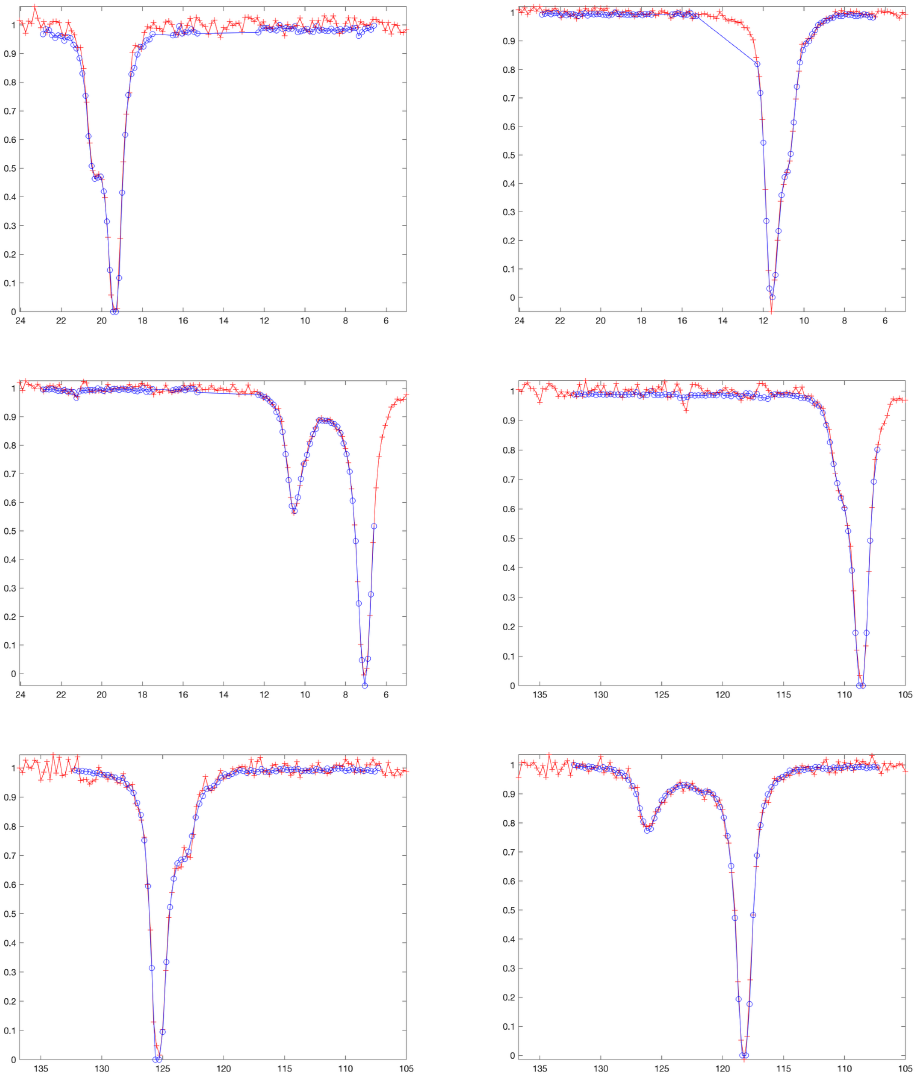
Comparison of representative CEST profiles obtained using ACCEST (red) or conventional CEST (blue). (a–c) ^13^C methyl data, (d–f) ^15^N data. CEST profiles are shown for residues (a) Leu55CD2, (b) Ile50CD, (c) Ile28CD, (d) Gly48, (e) Leu7, (f) Glu5. The number of scans was 8 for ^13^C- and ^15^N-ACCEST and ^15^N-CEST, and 4 for ^13^C-CEST.

Assuming similar signal-to-noise ratios (S/N), the ACCEST experiment reduces the total experiment time by a factor of *N*_*ω*1_/2, where *N*_*ω*1_ is the number of *ω*_1_ frequencies in the conventional CEST profile or the number of spectra recorded using D-CEST.^13, 14^ Whenever it is desirable to record multiple CEST profiles with different *B*_1_ field strengths, the total experiment time of conventional CEST or D-CEST essentially scales with the number of *B*_1_ field strengths. By contrast, ACCEST only requires one additional spectrum for each additional *B*_1_ field strength. In this case, the reduction in total experiment time is thus (*N*_ω1_ *N*_B1_)/(1+*N*_B1_), where *N*_B1_ is the number of *B*_1_ field strengths. Thus, ACCEST offers time savings of at least a factor of *N*_ω1_/2 and approaches *N*_ω1_ in the limit of large *N*_B1_. In practice, the effective time saving will be reduced by a factor of 2–3 as a greater number of scans may be needed in the ACCEST experiment to account for the inherently lower S/N of the ACCEST data (see Fig. 2d and 3).

The resolution of the CEST profiles acquired using ACCEST was 15.2 Hz/point and 22.6 Hz/point in the ^15^N and ^13^C experiments, respectively, whereas using conventional CEST the corresponding figures were 18.2 Hz/point (^15^N) and 22.6 Hz/point (^13^C). As indicated by the excellent agreement between ACCEST and CEST (Fig. 3), the difference in digital resolution of the CEST profiles have little impact on the quality of the data.

## METHODS

### Sample preparation

1 mM ^15^N-^13^CH_3_-ILV-^12^C^2^H isotope labeled A39V/N53P/V55L Fyn SH3 domain dissolved in H_2_O/D_2_O 90/10, 50 mM sodium phosphate, 2 mM EDTA, pH 7.0, and a small amount of NaN_3_. The ^15^NH and ^13^CH3 chemical shift assignments of A39V/N53P/V55L Fyn SH3 were obtained from the Biological Magnetic Resonance Data Bank, entry 17149.^22^

### NMR experiments

All experiments were acquired at 10 °C on a Bruker AVANCE NEO 600 MHz spectrometer equipped with a 5 mm QCI H&F/P/C-N-D CryoProbe and running Topspin 4.5.0.

The ^15^N ACCEST experiment was acquired with SW1 = 6211 Hz (oversampled by a factor of 3); *t*_1,max_ = 20.6 ms; number of *ω*_1_ frequencies, *N*_*ν*1_ = number of *t*_1_ values = TD1/2 = 128, where *ω*_1_ is moved from 105.0 ppm to 136.75 ppm with a step size of 0.25 ppm, using a *B*_1_-field of 15 Hz and duration 300 ms; recycle delay D1 = 2 s; number of scans NS = 8. The ^13^C methyl ACCEST experiment was acquired with SW1 = 10,000 Hz (oversampled by a factor of 3); *t*_1,max_ = 12.8 ms; *N*_*ν*1_ = TD1/2 = 128, where *ω*_1_ is moved from ppm to 20.05 ppm with a step size of 0.15 ppm, using a *B*_1_-field of 25 Hz and duration 300 ms; D1 = 2 s; NS = 8.

The conventional CEST experiments were acquired using the following parameter values. ^15^N: pulse sequence hsqc_cest_etf3gpsitc3d from the Bruker library; SW1 = 2070 Hz; *t*_1,max_ = 30.9 ms; *N*_*ν*1_ = 85, including a reference experiment without *B*_1_ irradiation, using a step size of 0.3 ppm and a *B*_1_- field of 15 Hz and duration 300 ms; TD1/2 = 64; D1 = 2 s; NS = 8. ^13^C methyl: pulse sequence mehsqccest3d from the Bruker library; SW1 = 3322 Hz; *t*_1,max_ = 19.3 ms; *N*_*ν*1_ = 86, including a reference experiment without *B*_1_ irradiation; TD1/2 = 64; D1 = 2 s; NS = 4.

The total time for each of the ^13^C methyl and ^15^N ACCEST experiments was 3 hours, including the reference experiment, whereas the total time for the conventional experiments were 60 hours for ^15^N and 30 hours for ^13^C methyl.

### Data processing

We processed the direct dimension (*t*_2_) using nmrPipe,^23^ employing squared cosine apodization and zero-filling prior to Fourier transformation. To generate CEST profiles from the ACCEST and reference spectra, we used the in-house software package (Dynamic Chemical Spectroscopy, DCS) as a front-end to the signal processing algorithm DSURE.^19, 21^

## ASSOCIATED CONTENT

### Supporting Information

Pulse sequence for the ^15^N-ACCEST experiment and detailed description of the data analysis workflow (PDF). The Supporting Information is available free of charge on the ACS Publications website.

## AUTHOR INFORMATION

### Author Contributions

GC and MA conceived the ACCEST approach and designed the data processing protocol. GC implemented the pulse sequence, recorded data, and wrote software implementing the processing protocol. TH produced the protein NMR samples. The manuscript was written by MA and GC with contributions from TH. All authors have given approval to the final version of the manuscript.

### Funding Sources

This research was supported by the Swedish Research Council (2021-05591), the Knut and Alice Wallenberg Foundation (2022.0085) and the European Union (ERC, DynaPLIX, SyG-2022 101071843). Views and opinions expressed are however those of the authors only and do not necessarily reflect those of the European Union or the European Research Council. Neither the European Union nor the granting authority can be held responsible for them.

## ACKNOWLEDGMENT

We thank Andreas Jakobsson for helpful discussions.

## ABBREVIATIONS

ACCEST: accordion-CEST
CEST: chemical exchange saturation transfer
HSQC: heteronuclear single-quantum coherence

## Supplementary Information

### Processing Procedure: Reconstruction of CEST Profiles

We first use a generic CEST profile to generate estimates of the CEST profiles for all spins *k*, starting from the spin *k* with lowest amplitude. The generic CEST profile can be obtained from an isolated resonance in the spectrum which does not exchange with a minor conformer. The estimated profiles are successively subtracted from Δ*S*^r^_j_(*t*_1_). The profile obtained for a given spin *j* is saved and used later in analyses of all neighboring *t*_1_ columns where this resonance is present. Next, a *t*_1_ column where one of the spins *k* is at maximum intensity is analyzed to obtain a better estimate of the CEST profile for this spin. Residual signals from other spins in the column are subtracted, using either the saved CEST profile if it exists, or the generic profile. Again, the result is saved and later used for this spin in subsequent analyses. This is repeated until all spins *k* present in the *t*_1_ column have updated CEST profiles. The procedure is then repeated, starting from spin *j*, until convergence, which typically requires only 2–4 interations. SI Figure 1 outlines a block diagram describing the reconstruction procedure.

**SI Figure 1.**
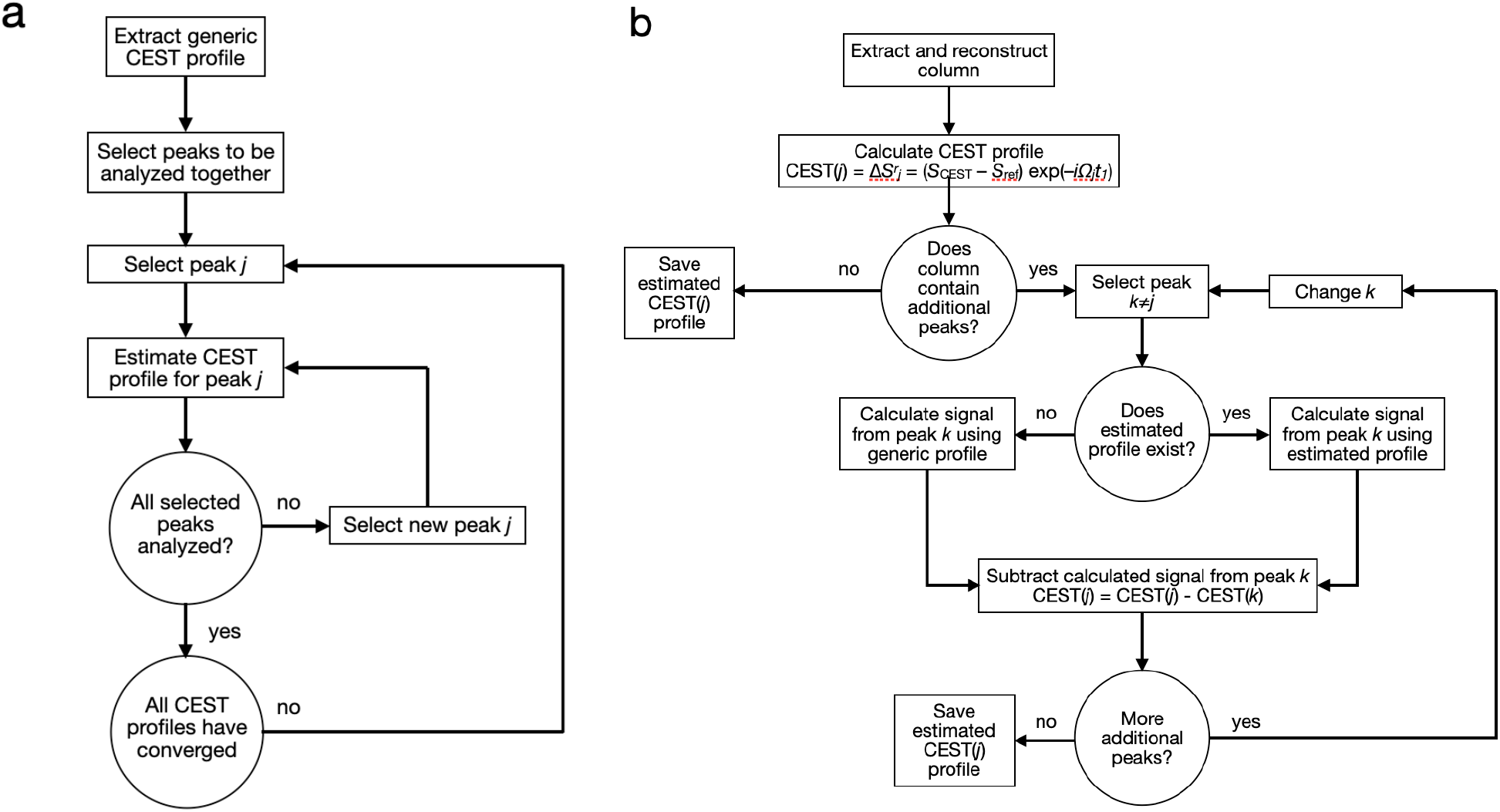
Block diagrams showing the workflows used to obtain CEST profiles from ACCEST data. (a) Workflow for iterative analysis of a group of resonance peaks appearing in each other’s *t*_1_ interferograms. (b) Workflow for estimation of CEST profile for a single peak, correcting for additional resonances in the *t*_1_ interferogram.

## Notes

### Competing Interest Statement

The authors have declared no competing interest.

